# Time-adaptive Unsupervised Auditory Attention Decoding Using EEG-based Stimulus Reconstruction

**DOI:** 10.1101/2022.01.07.475386

**Authors:** Simon Geirnaert, Tom Francart, Alexander Bertrand

**Author notes:** (Corresponding author: Simon Geirnaert.);.

## Abstract

The goal of auditory attention decoding (AAD) is to determine to which speaker out of multiple competing speakers a listener is attending based on the brain signals recorded via, e.g., electroencephalography (EEG). AAD algorithms are a fundamental building block of so-called neuro-steered hearing devices that would allow identifying the speaker that should be amplified based on the brain activity. A common approach is to train a subject-specific stimulus decoder that reconstructs the amplitude envelope of the attended speech signal. However, training this decoder requires a dedicated ‘ground-truth’ EEG recording of the subject under test, during which the attended speaker is known. Furthermore, this decoder remains fixed during operation and can thus not adapt to changing conditions and situations. Therefore, we propose an online time-adaptive unsupervised stimulus reconstruction method that continuously and automatically adapts over time when new EEG and audio data are streaming in. The adaptive decoder does not require ground-truth attention labels obtained from a training session with the end-user and instead can be initialized with a generic subject-independent decoder or even completely random values. We propose two different implementations: a sliding window and recursive implementation, which we extensively validate on three independent datasets based on multiple performance metrics. We show that the proposed time-adaptive unsupervised decoder outperforms a time-invariant supervised decoder, representing an important step toward practically applicable AAD algorithms for neuro-steered hearing devices.

## I. INTRODUCTION

The auditory attention decoding (AAD) task consists of determining to which speaker out of multiple simultaneously talking speakers a listener wants to attend. Such AAD technology could be employed in hearing aids, cochlear implants, or other so-called ‘hearables’, to inform beamformers and noise suppression algorithms about which speaker to enhance and which other speakers to treat as background noise and thus to suppress. Heuristics such as look direction can be used to make an informed guess, yet such indirect metrics often fail as they do not always sufficiently correlate with the actual auditory attention. Ideally, the auditory attention can be directly decoded at the source, i.e., the brain. It has been extensively shown that the auditory attention information is encoded in brain signals [1]–[3], which can be recorded, for example, using electroencephalography (EEG). As such, EEG-based AAD technology could contribute to so-called ‘neuro-steered’ hearing devices [4], [5].

The predominant paradigm for AAD is stimulus reconstruction [6]. A stimulus decoder, consisting of a linear spatio-temporal filter, is applied to multi-channel EEG data of the listener to reconstruct the speech envelope of the attended speaker. By computing the correlation between this reconstructed speech envelope and the original speech envelopes of the different competing speakers, the attended speech signal and corresponding speaker can be identified [5]–[7]. However, such a stimulus decoder is traditionally trained in a supervised way, i.e., it assumes the attended speaker is known during the training phase of the decoder. Such ground-truth labels can be obtained by instructing the subject to attend to a specific speaker during a dedicated EEG recording session.

In [8], we proposed an unsupervised algorithm to train a subject-specific stimulus decoder without the need for ground-truth labels. Consequently, the first issue with the traditional stimulus reconstruction method, i.e., the need for acquiring labeled data during a dedicated training session, is resolved while approximating the performance of a fully supervised subject-specific decoder [8]. However, this unsupervised decoder is still trained in batch on a large amount of (unlabeled) training data and then remains fixed during operation. Such fixed decoders do not adapt to long-term signal changes due to changing conditions and situations (e.g., non-stationarities in the neural activity, changing electrode-skin contact impedances, shifting or loosening electrodes). Therefore, in this paper, we modify and extend the algorithm proposed in [8] such that the decoder adapts over time in an unsupervised manner. The resulting decoder does not require a dedicated training session and can automatically adapt to new incoming non-stationary EEG data from the end-user and thus serves as one of the first practical plug-and-play AAD algorithms for neuro-steered hearing devices.

In Section II, we review the (unsupervised) stimulus reconstruction algorithm for AAD. In Section III, we then present and explain the proposed time-adaptive updating schemes. These implementations are investigated and the hyperparameter choices are validated in Section IV. The proposed time-adaptive unsupervised decoder is then tested and compared to the fixed (time-invariant) supervised decoder in a time-adaptive context in Section V, by simulating a scenario where electrodes are disconnected and applying the decoders on a dataset recorded across multiple recording days.

## II. (UN)SUPERVISED STIMULUS RECONSTRUCTION FOR AAD

### A. Review of stimulus reconstruction

Consider a *C*-channel EEG signal of which the *c*^th^ channel is denoted by *x*_*c*_(*t*), with *t* the time sample index. In the linear stimulus reconstruction paradigm, a spatio-temporal filter or decoder *d*_*c*_(*l*) is applied to this *C*-channel EEG signal to reconstruct the speech envelope of the attended speaker *s*_a_(*t*) [5]–[7]:

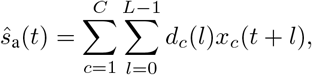

with the channel index *c* ranging from 1 to *C* (spatial combination of *C* channels) and the time lag index *l* ranging from 0 to *L* – 1 (temporal integration over *L* time samples). This filter is an anti-causal filter, as *L post-stimulus* time lags are used to reconstruct the attended speech envelope from the EEG signal. To identify the attended speaker, the reconstructed envelope 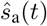 from the EEG is compared with original speech envelopes^1^ *s*_1_(*t*) and *s*_2_(*t*) of the two simultaneously talking speakers through the Pearson correlation coefficient. For the sake of an easy exposition but without loss of generality, we here assume only two competing speakers, although all presented algorithms and procedures can be extended to more speakers.

In the remainder of the paper, we will adopt a matrix-vector notation, in which the decoder is written as **d** = [*d*_1_ (0) *d*_1_ (1)…*d*_1_ (*L* – 1) *d*_2_ (0)…*d*_*C*_ (*L* – 1)]^T^ ∈ ℝ^*CL*^. Assume (for now) the availability of *K* training segments of *T* time samples, where the available training information is described as 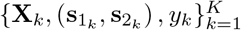, containing an EEG data **X**_*k*_ matrix collecting all *T* time samples in training segment *k*; a rigorous definition is given in (3)), speech envelopes 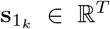 and 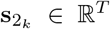 (similarly), and attention labels *y*_*k*_ ∈ {1, 2}, indicating which speech envelope (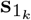 or 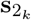) is the attended one 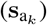 (assuming constant attention across the whole segment). For each training segment *k*, the attended speech envelope is determined as

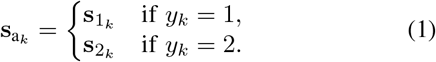

The decoder is then trained by minimizing the squared error between the actual attended and reconstructed speech envelope across all training segments:

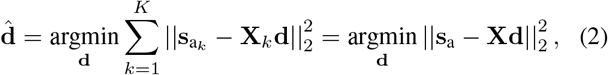

with 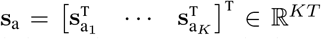 the concatenated actual attended speech envelope and where the block Hankel matrix **X** ∈ ℝ^*KT* × *CL*^ represents the concatenated time-lagged *C*-channel EEG with *L* time lags:

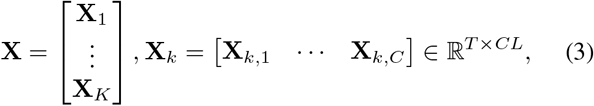

with **X**_*k,c*_ ∈ ℝ^*T* × *L*^ a Hankel matrix containing the time-lagged EEG data of the *k*^th^ training segment and *c*^th^ EEG channel:

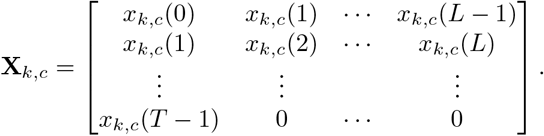

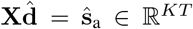 then represents the reconstructed speech envelope over all training segments. The solution of (2) is found by solving the normal equations:

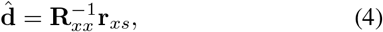

with

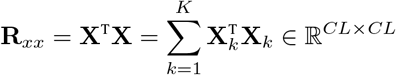

the estimated EEG autocorrelation matrix and

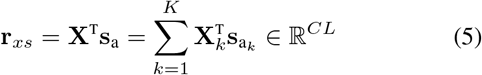

the estimated crosscorrelation vector between the EEG and the attended speech envelope. It is important to notice that only in (5) we need the attention labels *y*_*k*_ to select the attended speech envelope 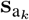 in segment *k* (see (1)). We use shrinkage to regularize the estimated autocorrelation matrix:

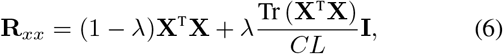

with **I** ∈ ℝ^*CL*×*CL*^ the identity matrix and where the shrinkage parameter 0 ≤ *λ* ≤ 1 is analytically determined [10], [11]:

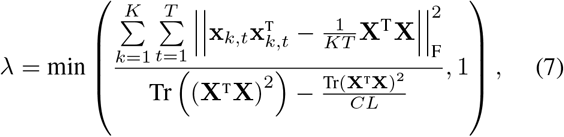

with 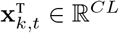 the *t*^th^ row of the matrix **X**_*k*_.

Given the estimated decoder 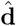 and *T*_test_ time samples of a new EEG segment 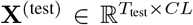 of a subject listening to one out of two competing speakers with speech envelopes 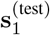 and 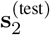, a decision about the auditory attention of the listener can be made by:

1. reconstructing the attended speech envelope from the EEG: 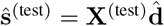 and
2. computing the Pearson correlation coefficients 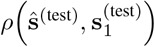 and 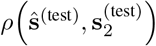 between this reconstructed speech envelope and the original speech envelopes. The speaker corresponding to the highest correlation coefficient is identified as the attended speaker.

For stimulus reconstruction, there is an important trade-off between the accuracy of the decision and the decision segment length *T*_test_, i.e., the number of time samples used to make a decision [5], [12]. A longer decision segment leads to more accurate estimates of the Pearson correlation coefficients, thereby improving accuracy on the AAD decisions. However, this comes with the drawback of a poorer time resolution at which the AAD decisions are made due to the longer decision segment length.

### B. Unsupervised stimulus reconstruction

Let us now assume that the attention labels 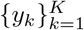 are not known, even during training, meaning that we do not know to which of the speakers the subject is attending when designing our decoder (i.e., using (1) has become impossible). If we indeed can train the decoder without these labels, this would avoid the need for a dedicated training session during which the subject is instructed to attend to a specific speaker in order to collect ground-truth data. As a result, performing AAD becomes an *unsupervised* classification problem. The absence of labels is a roadblock in the computation of the crosscorrelation vector in (5), which requires the use of the correct speech envelope in its calculation (using the unattended envelope in (5) would result in a decoder that emphasizes the wrong speaker).

In [8], we proposed an unsupervised batch-training procedure on an unlabeled dataset 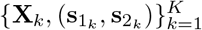 of *K* segments (i.e., attention labels 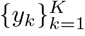 are unavailable/unknown). The main idea is to iteratively retrain a decoder by using labels that are predicted by the decoder from the previous iteration. A short description of this iteretive procedure is as follows. We start with an initial decoder 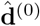 at iteration 0, which can be, e.g., a pre-computed subject-independent decoder or even a decoder with random entries. First, the EEG autocorrelation matrix **R**_*xx*_ is estimated as in (6), which does not require any attention labels. The iterative prediction of the labels and updating of the decoder then comprises the following steps:

1. given a decoder 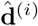 that is available after the *i*^th^ iteration, apply it on each EEG segment **X**_*k*_ to reconstruct the attended speech envelope:

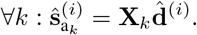
2. Per segment, correlate the reconstructed speech envelope 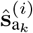 with both speech envelopes 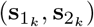 to predict the attended speaker. As before, the first speaker is idenatsifitehde atten d one if the mple Pearson correlation coefficient 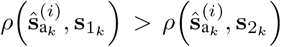 and vice versa. The predicted attended envelope is denoted as 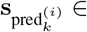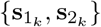.
3. Using the EEG segments and corresponding speech envelopes of the predicted attended speaker 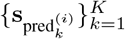, the crosscorrelation vector can be computed/updated as in (5). It is crucial to use the *original* speech envelope (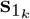 or 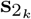) and not the envelope 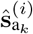 that was reconstructed from the EEG. Given the new crosscorrelation vector, the decoder 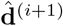 can be updated as in (4). Then return to step (1) and iterate until convergence.

This iterative unsupervised predicting of labels could potentially inject incorrect labels and thus incorrect data in the estimation of the decoder, which could in principle lead to a downward spiral of incorrect updating and thus to a badly-performing decoder. Remarkably, it was shown in [8] that a *self-leveraging effect* occurs in this batch-mode iterative updating where the new decoder outperforms the previous decoder, despite the presence of labeling errors, resulting in an upward instead of a downward spiral. This happens even in the case where the initial decoder is initialized with random values. This unsupervised subject-specific decoder outperformed a supervised subject-independent decoder (i.e., trained on data from other subjects than the one under test) and even closely approximated the performance of a supervised subject-specific decoder [8].

## III. TIME-ADAPTIVE UNSUPERVISED STIMULUS RECONSTRUCTION FOR AAD

The unsupervised training procedure of Section II-B assumes the availability of multiple data segments at once (i.e., batch-training). The batch computation is inherent to the procedure; once a new decoder is computed, *all* labels in the recording are repredicted to improve the next decoder. After the unsupervised batch training, the final decoder is fixed and applied to unseen data from the subject under test. However, such a pre-trained time-invariant decoder does not adapt to non-stationarities due to changing conditions and situations and may thus perform suboptimally. Here, we propose a time-adaptive realization of such an unsupervised AAD decoder, i.e., a decoder that adapts itself over time when EEG and audio data (processed in envelopes) are continuously streaming in.

Assume some initial decoder. Data segments of *T*_ud_ samples of EEG and audio data start streaming in. At a certain point in time, assume the *k*^th^ segment of EEG data 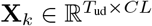 (see (3)) and corresponding segments of the speech envelopes of the two competing speakers 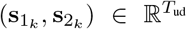 become available. There is no information available about which speaker is the attended or unattended one in this segment. The goal is now to update the decoder in an unsupervised manner based on the newly available information, to which end we will propose and compare two approaches (Section III-A and III-B).

In this time-adaptive procedure, it is important to distinguish the updating segment length *T*_ud_ from the decision segment length *T*_test_. The former, equivalent to the segment length *T* in the previous sections, corresponds to the length of the segments on which the prediction of the labels for the updating/training and the updating/training itself is performed. The latter corresponds to the length of the segments on which AAD decisions are made to in the end steer the enhancement algorithm in the hearing device. This decision segment length is, therefore, much more sensitive to speed (e.g., because of switches in auditory attention [12]) than the updating segment length, as there can be some delay allowed in updating the decoder. Therefore, the updating segment length is typically larger than the decision segment length *T*_ud_ ≥ *T*_test_, i.e., within each updating segment, multiple AAD decisions are made.

In the time-adaptive sliding window approach Section III-A), the aforementioned batch-mode procedure is mimicked (i.e., including repredictions of the labels of previous segments) but over a finite time horizon that is implemented as a sliding window. In Section III-B, we propose an alternative time-adaptive approach that does *not* recompute previously predicted labels and, therefore, can be implemented recursively. This is much more attractive from a computational and memory usage point of view. Both approaches to update the decoder are explained in more detail in the following sections.

### A. Sliding window implementation

In the sliding window implementation (Figure 1), a pool of *K* data segments of updating segment length *T*_ud_ is permanently kept in memory using a first in, first out (FIFO) principle. When a new data segment 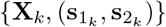 becomes available, the oldest data segment is discarded and the pool is updated with the newest one, resulting in the new pool 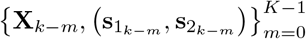. The stimulus decoder is then updated similarly to the batch-mode implementation explained in [8] and Section II-B, but on the finite pool of *K* segments.

**Figure 1:**
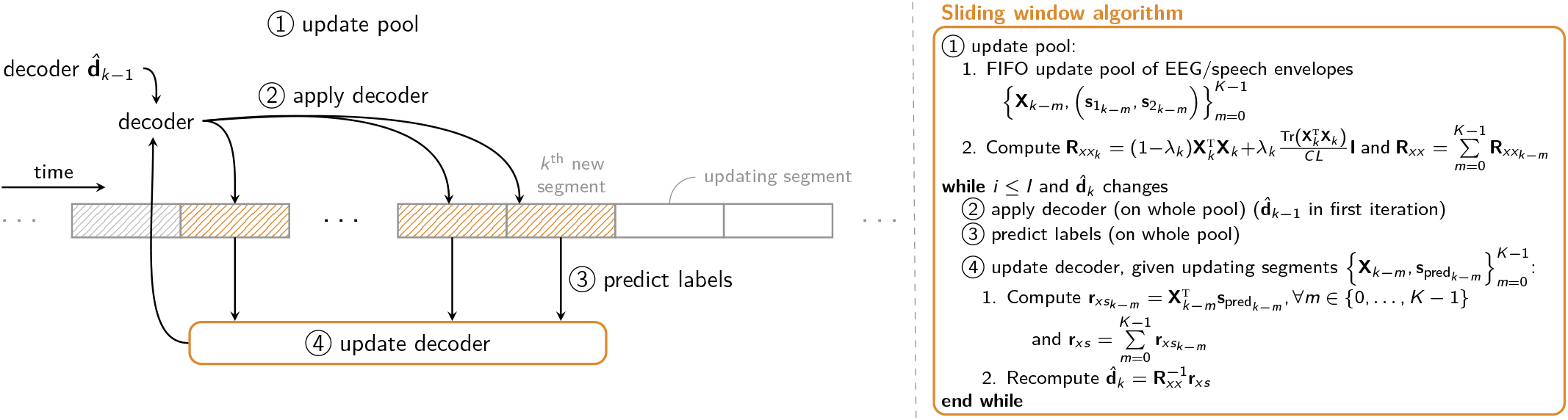
The time-adaptive unsupervised sliding window scheme and algorithm to update a stimulus decoder, with repredictions of the labels on previous segments.

First, the EEG autocorrelation matrix of the new segment is computed similarly to (6):

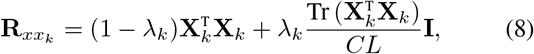

with the regularization parameter *λ*_*k*_ recomputed per new segment *k* using (7). The aggregated autocorrelation matrix across the whole pool of *K* segments can then be updated/recomputed:

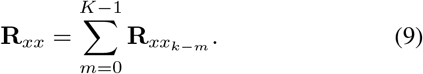

The autocorrelation matrices of the previous segments can be recomputed or stored and retrieved from previous computations. We found empirically that better results are obtained when regularizing the new autocorrelation matrix (8) *before* being stored and combined in (9), instead of regularizing the combined autocorrelation matrix (9), i.e., *after* combining the autocorrelation matrices from the different segments.

Using the decoder 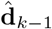 from the previous step, the iterative procedure of predicting labels, updating the crosscorrelation vector(s), and decoder on the pool of *K* segments 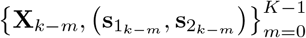 can be initiated. Given the persegment predicted attended speaker 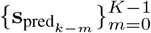 (initially obtained using 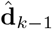), the crosscorrelation vectors can be updated as in (5):

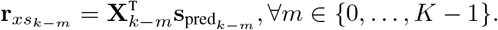

The aggregated crosscorrelation vector and corresponding decoder can then be computed as:

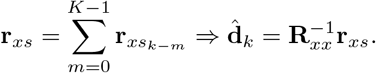

This predict-and-update procedure is then iterated *I* times on the same pool of *K* segments. Based on [8], we choose *I* = 5, after which the iterative batch updating procedure is generally observed to have converged to a final decoder [8] (see also Figure 6). Given that we iterate over this pool of *K* segments, this approach can only be implemented in a sliding window manner and not recursively. Lastly, the pool size parameter *K* represents an important trade-off between accuracy, adaptivity, and computational complexity and memory usage. A longer sliding window (i.e., larger *K*) means that more data is available to compute the decoder, resulting in a better approximation of the batch-mode decoder but also resulting in a lower adaptivity and higher memory requirements (see also Section IV).

**Figure 2:**
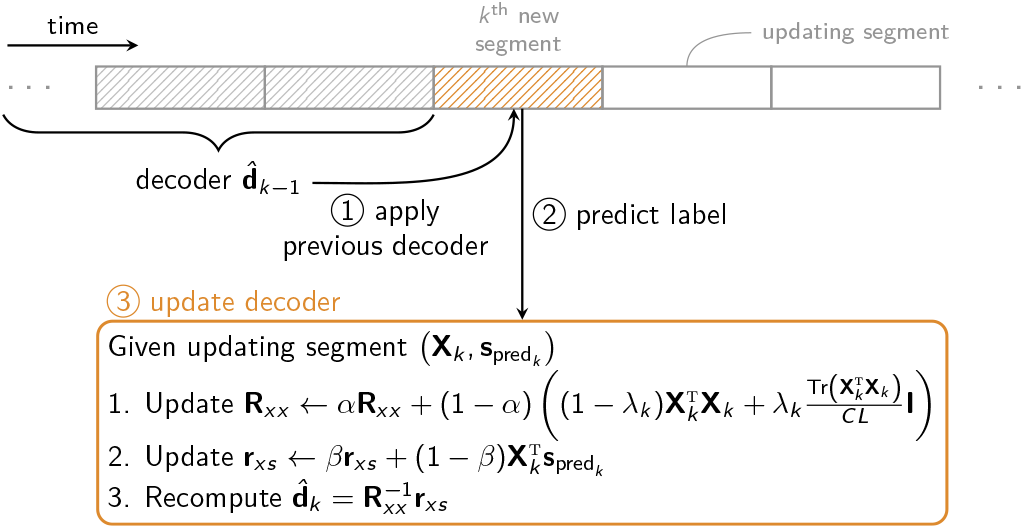
The time-adaptive unsupervised recursive predict-and-update scheme to update a stimulus decoder.

**Figure 3:**
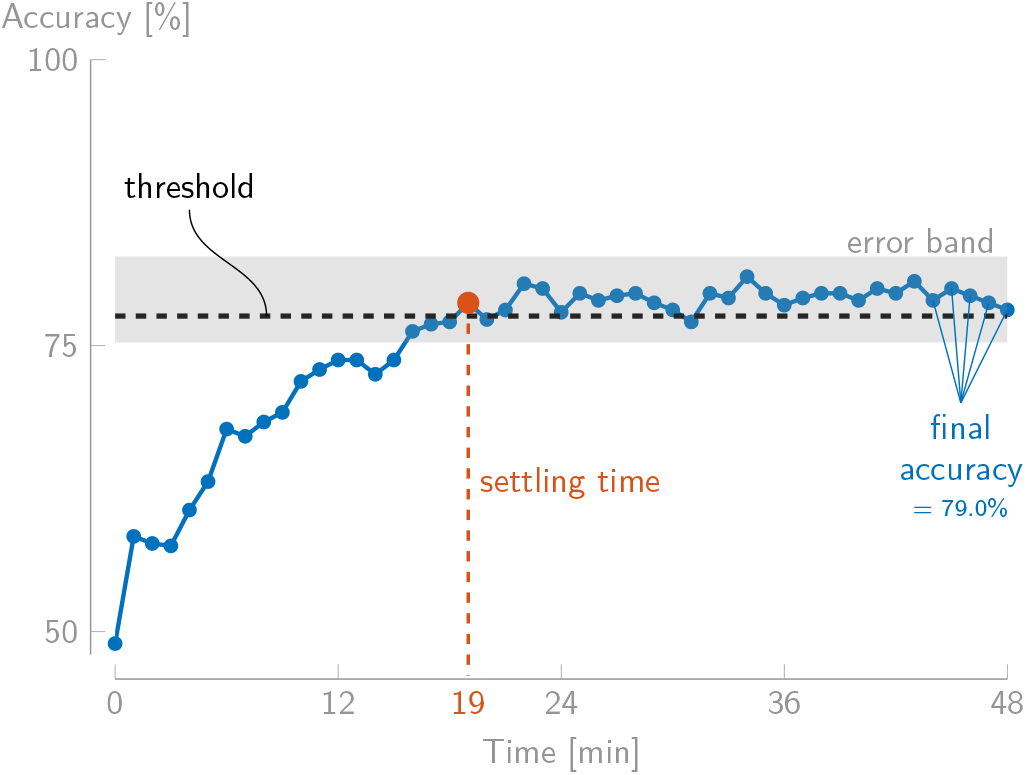
Illustration of the adaptation curve and performance metrics for a representative subject (Subject 4) of Dataset I, starting from a random initial decoder and updating every 60 s, for the recursive implementation with *α* = *β* = 0.9 (average across ten runs).

**Figure 4:**
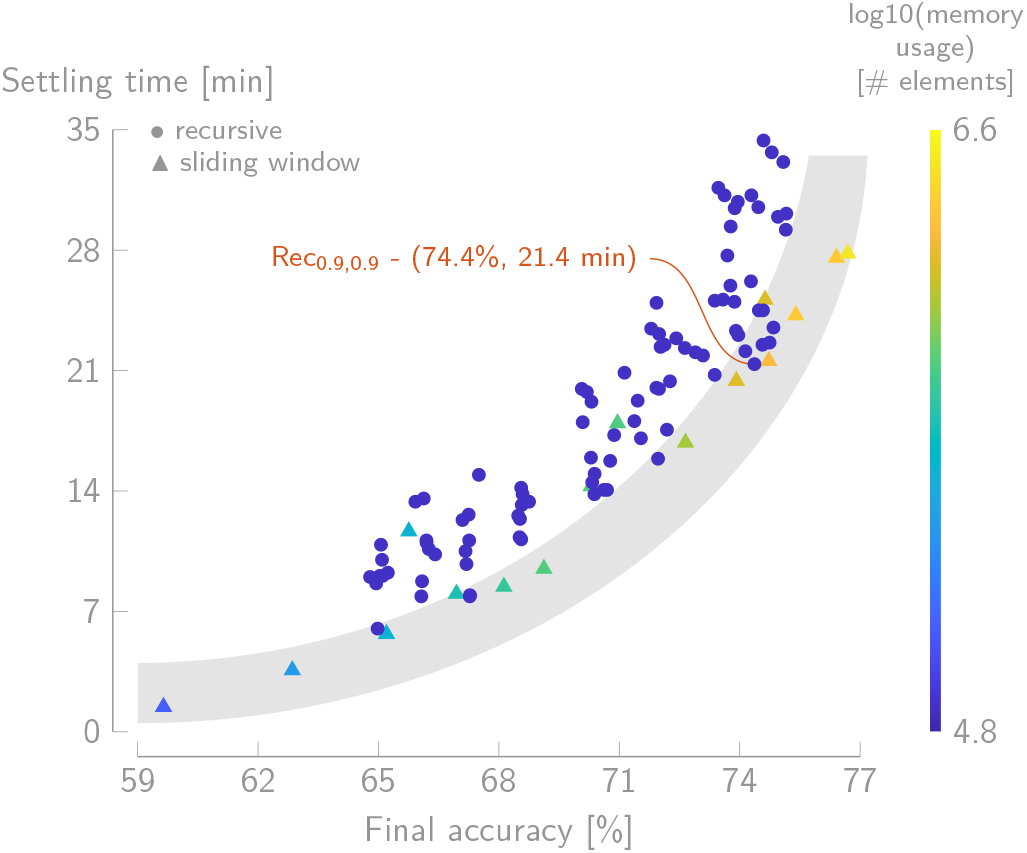
Average settling time vs. final accuracy for different parameter settings across the 16 subjects of Dataset I and ten random permutations per subject. The shaded area highlights the points that are close to or in the Pareto front. The indicated recursive algorithm with *α* = *β* = 0.9 gives one of the best trade-offs between final accuracy, adaptivity, and memory usage across the evaluated settings.

**Figure 5:**
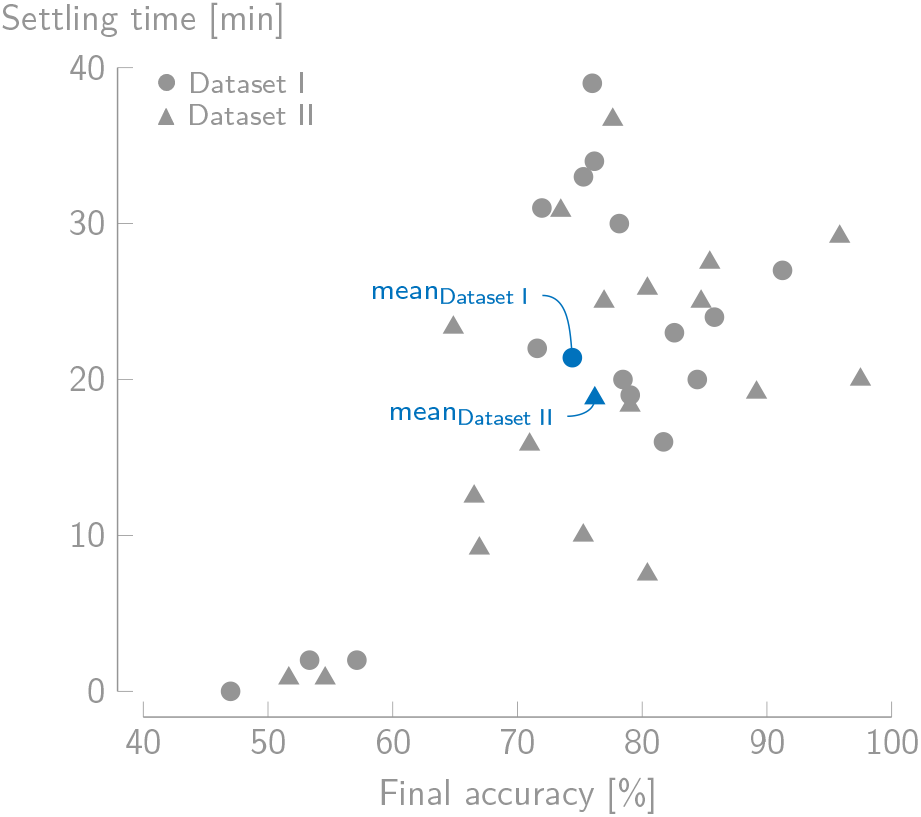
Individual settling time vs. final accuracy per subject of Dataset I and II (average across runs) for the chosen recursive implementation with *α* = *β* = 0.9*, T*_ud_ = 60 s (Dataset I) and *α* = *β* = 0.916*, T*_ud_ = 50 s (Dataset II).

**Figure 6:**
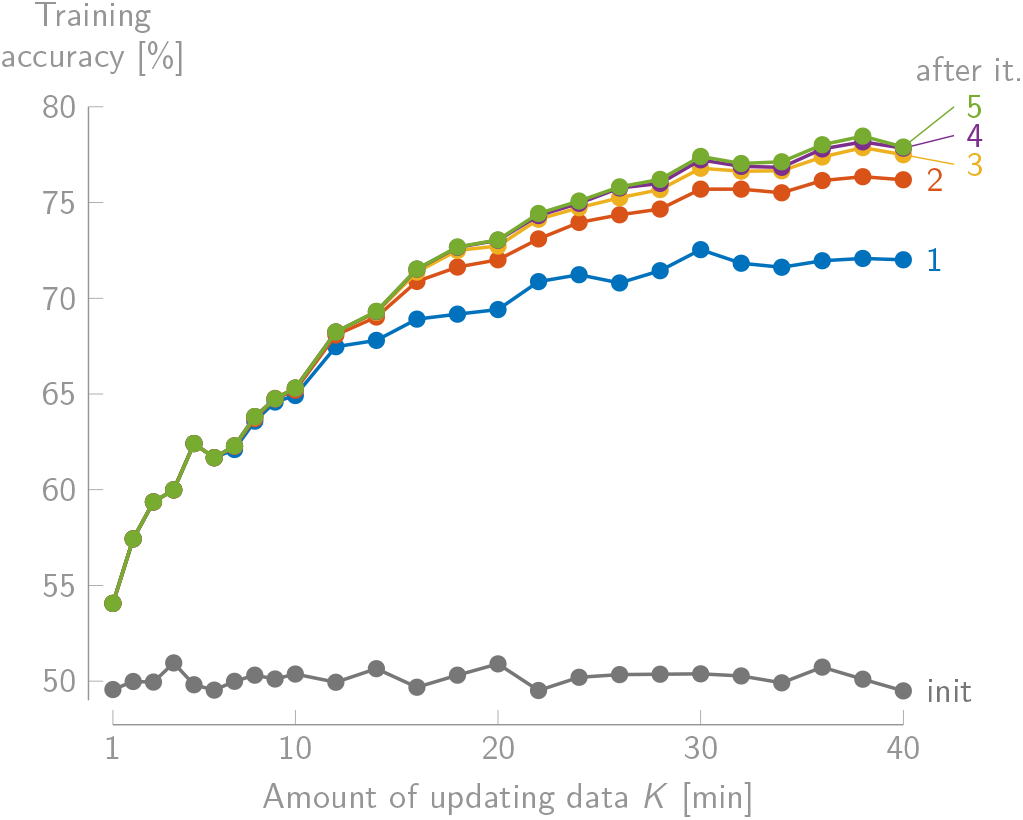
The training accuracy on 30 s decision segments for the batch-mode unsupervised iterative updating procedure as a function of the amount of updating data, for different numbers of relabeling iterations (‘init’ refers to the accuracy when no iterations are performed; average across all subjects of Dataset I and random permutations). Given that the updating segment length *T*_ud_ = 60 s, the amount of updating data corresponds to pool size *K* (in number of minutes).

### B. Recursive implementation

As an alternative to the sliding window implementation, we propose a single-shot predict-and-update scheme (Figure 2). As opposed to the sliding window approach, the labels of previous updating segments are not repredicted, enabling a recursive implementation that is much more efficient from a computational and memory usage point of view. In this recursive implementation, the decoder resulting from the previous update 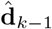 is applied to the new segment 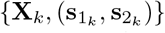 of length *T*_ud_ to predict the label of this new segment, resulting in the predicted attended envelope 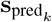. To update the decoder, a regularized autocorrelation matrix is computed based on the new *k*^th^ segment as in (8), while the predicted attended envelope is used to compute a crosscorrelation vector as in (5):

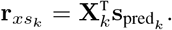

This new autocorrelation matrix crosscorrelation vector 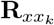 and crosscorrelation vector 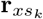 can then be combined with the autocorrelation matrix **R**_*xx*_ and crosscorrelation vector **r**_*xs*_ integrating all previous information to update the decoder:

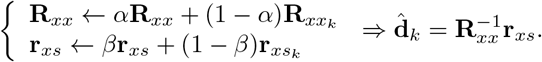

The influence of the weighting parameters *α* and *β* will be empirically evaluated in Section IV.

Unlike the sliding window implementation, which uniformly weighs the *K* past segments in the new decoder, this recursive algorithm implements an exponential weighting across all past segments. This exponential weighting could be advantageous, especially in an adaptive context, as the more relevant closest (past) segments have higher weights than those further in the past. One can choose the weighting parameters *α* and *β* such that the center of mass of the exponential weighting is the same as of the sliding window approach [13]:

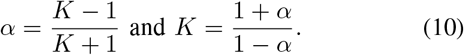

A possible drawback of this recursive implementation is that previous labels are not repredicted, and that one can not apply multiple iterations over a pool of segments as in the sliding window version. This could lead to slower convergence or poorer accuracies. However, the upside is that the procedure is much easier to implement and much more efficient in terms of memory and computation resources (see Section III-C). In Section IV, we will demonstrate that the impact on convergence speed and accuracy is negligible.

### C. Memory usage

#### 1) Sliding window implementation

For the sliding window implementation, at least the pool of *K* EEG segments (*K* × *T*_ud_ × *C*) and 2*K* speech envelopes (2*K* × *T*_ud_) need to be permanently stored in memory. One does not need to store the *L* different time lags, as these can always be generated from the original EEG data. Furthermore, it is not possible to simply store the autocorrelation matrices and crosscorrelation vectors of the previous EEG segments - which would require less memory usage - as for the repredictions of the labels, the decoder has to be applied to the original EEG and speech envelope data. This leads to the following memory usage:

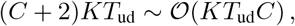

where generally *T*_ud_ >> *K* or *C*. As the storage of the sample dimension *T*_ud_ is required, this is generally a very high memory usage.

#### 2) Recursive implementation

The recursive implementation minimally requires the permanent storage of one auto-correlation matrix **R**_*xx*_ (built from 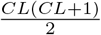 elements due to symmetry) and one crosscorrelation vector **r**_*xs*_ (built from *CL* elements). This leads to the following memory usage:

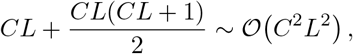

which is, as expected, much less than the sliding window approach.

To better appreciate the differences, consider the following practical realistic example with *C* = 24 EEG channels, *L* = 6 time lags, pool size *K* = 19, and an updating segment length *T*_ud_ equivalent to 1200 samples (corresponding to 60 s when the EEG and speech envelopes are downsampled to 20 Hz). The sliding window approach then requires the storage of 592 800 elements, which is more than 50 times more than for the recursive implementation, which requires to only story 10 584 elements.

## IV. VALIDATION AND COMPARISON

We test both versions of the time-adaptive unsupervised stimulus reconstruction algorithm of Section III for different hyperparameter settings (i.e., the updating segment length *T*_ud_, pool size *K* in the sliding window implementation, and exponential forgetting factors *α, β* in the recursive implementation). In all experiments, we start from a different *fully random* initial decoder, generating the first prediction(s). In the recursive implementation, the initial autocorrelation matrix and crosscorrelation vector are initialized with all zeros. We compare the sliding window and recursive implementation and select a set of hyperparameters on one dataset and validate the chosen algorithm on a second one. These datasets are introduced in Section IV-A, while the performance metrics are described in Section IV-B. The experiments and results are discussed in Section IV-C, with a more detailed discussion on the effect of repredictions of the labels in the sliding window approach in Section IV-D. The final settings are validated on the second dataset in Section IV-E.

### A. Data and preprocessing

#### 1) AAD datasets

The first AAD dataset (Dataset I) is from [9] and contains the EEG (64-channel BioSemi ActiveTwo system, standard 10-20 layout, 8196 Hz sample rate) and audio data of 16 normal-hearing subjects participating in an AAD experiment, where the subjects were instructed to listen to one of two competing speakers located at ±90° azimuth direction (in dichotic and head-related transfer function (HRTF)-filtered listening conditions). Per subject, eight stories of 6 min and 12 repetitions of 2 min of those stories are presented, resulting in 72 min data per subject. The attention was balanced across left and right attended and listening conditions. During the experiments, the eyes of the participants were open. This dataset is available online [14], and we refer to [9] for more details.

The second AAD dataset (Dataset II) is from [15] and will act as an independent validation dataset. It contains the EEG (64-channel BioSemi ActiveTwo system, standard 10-20 layout, 512 Hz sample rate) and audio data of 18 normal-hearing subjects in a similar AAD experiment with two competing speakers, located at ±60° azimuth direction (HRTF-filtered) and using different acoustic room properties. Per subject, 50 min of data (60 × 50 s trials) are available. The attention was again balanced across listening directions and conditions. The eye gaze of the participants was fixed to a crosshair. This dataset is also available online [16], and we refer to [15] for more details.

#### 2) Preprocessing

To preprocess the EEG and audio data, we applied the same preprocessing steps as in [8], [9]. The speech signals are first filtered using a gammatone filterbank. Using a power-law operation with exponent 0.6, an envelope is computed for each subband signal. All subband envelopes are afterward summed to one envelope. Both EEG and speech envelopes are bandpass filtered between 1–9 Hz and down-sampled to 20 Hz (Dataset I)/32 Hz (Dataset II). In neither of the datasets, additional re-referencing or artifact rejection has been applied.

### B. Performance metrics

To evaluate a specific implementation with a specific set of hyperparameters, we use three performance metrics, quantifying the accuracy, adaptivity, and memory usage of each algorithm:

- *Final accuracy:* the final accuracy is defined as the average of the accuracies on the independent test set across the last 5 min of updating, i.e., after the adaptive decoder has had sufficient time to converge to a steady-state regime.
- *Settling time:* in a time-adaptive context, not only accuracy but also adaptivity or speed of adaptation is an important metric. Here, we quantify the adaptivity with the settling time, defined similarly as in control theory [17]. This settling time is defined as the point in time where the accuracy has reached a threshold for the first time *and* remains above the lower bound of a predefined error band for the remainder of the updating procedure. The threshold is defined as a convex combination of the final accuracy and the initial chance level performance (50%; before the updating procedure):

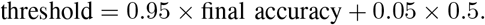 The error band, which allows taking the variability into account, is defined as:

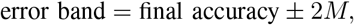

with *M* the difference between the maximum and minimum across the last 5 min accuracies.
- *Memory usage:* the memory usage, i.e., the number of elements that need to be permanently stored in memory, is computed as in Section III-C.

Figure 3 illustrates the final accuracy and settling time performance metrics for a representative subject and specific implementation (this figure is only meant to illustrate the updating procedure and the definitions of the performance metrics, and should not be viewed as a validation result).

### C. Hyperparameter selection

We test the different implementations of Section III for different hyperparameter settings on Dataset I. Per subject, we randomly permute the 6 min trials of the first 48 min and use those as the updating set, i.e., the data on which the time-adaptive unsupervised updating from a random initial decoder is performed. To track the accuracy of the updated decoder over time, after each update, we evaluate the decoder on the separate set of the last 24 min of repetition data, using *T*_test_ = 30 s decision segments to make a decision about the auditory attention (i.e., to compute the Pearson correlation coefficient with both speech envelopes). Per subject, we perform ten random permutations. For the decoder, we choose time lags up to 250 ms [6], [9], which corresponds to *L* = 6 for Dataset I and *L* = 9 for Dataset II (as both are sampled at different rates). We choose updating segment length *T*_ud_ = 60 s (different from decision segment length *T*_test_ = 30 s), i.e., we update the decoder every 60 s. As the performance of the stimulus decoder heavily depends on the amount of data available to make a decision [5], [12], similarly to [8], we choose *T*_ud_ as large as possible - without waiting too long to produce as reliable labels as possible. As explained in the introduction of Section III, we can afford such a longer delay in updating (as opposed to the decision segment length, explaining why we work on different time resolutions for the updating and testing). However, we do not take *T*_ud_ larger than 60 s, as the performance of the stimulus decoder starts to saturate above this segment length, and because this would require a too long sustained attention on the same speaker.

Figure 4 shows the average final accuracy, settling time, and memory usage for different settings of the sliding window and recursive implementations across the 16 subjects of Dataset I and ten random permutations. To compute the final accuracy per setting, 16 (number of subjects) ×10 (random permutations) ×5 (number of time points of updating; see Section IV-B) ×48 (number of 30 s decision segments in the 24 min test set) evaluations are thus performed. The sliding window implementation is evaluated for different updating segment lengths (*T*_ud_ ∈ {60, 30, 10 s}) and pool sizes (*K* ∈ {10, 20,…, 60}, except for *T*_ud_ = 60 s, where the maximum is *K* = 40). Only the results for *T*_ud_ = 60 s updating segments are shown, which are found to be superior to *T*_ud_ = 30 s and 10 s. Therefore, and for the clarity of Figure 4, the recursive implementation is only evaluated for *T*_ud_ = 60 s and *α, β* ranging independently from each other from 0.6 to 0.95 in steps of 0.05 and *α* = *β* ranging in more fine-grained steps of 0.005 from 0.8 to 0.95. As can be seen in Figure 4, the memory usage of the recursive implementation is the same for every setting (Section III-C2), while this is dependent on the pool size *K* and updating segment length *T*_ud_ for the sliding window implementation (Section III-C1). In general, there is a clear positive correlation between a higher final accuracy and a higher settling time, representing the trade-off between accuracy and adaptability of the decoder. The points in the shaded lower envelope area of the point cloud in Figure 4 represent the Pareto front, i.e., the settings that give the best trade-off between a high final accuracy and low settling time.

Surprisingly, the Pareto front of the recursive implementations (i.e., without repredictions of previous labels) seems to achieve very similar performances in terms of final accuracy and settling time as the sliding window implementations (i.e., with repredictions of previous labels). In Section IV-D, we will investigate more closely why these repredictions of previous labels seem to have such little effect. Moreover, the recursive implementation requires on average 16× less memory than the sliding window implementation (see Figure 4) and is computationally more efficient, making it the preferred implementation.

As indicated in Figure 4, one of the best choices across the evaluated settings is the recursive implementation with *α* = *β* = 0.9, resulting on average in 74.4% final accuracy (standard deviation 12.1%) after 21.4 min (st. dev. 11.7 min) (see Figure 5 for the per-subject performances). The latter implies that it takes about 20 min before the decoder has learned how to optimally decode the attended speaker starting from a random decoder. These settings are therefore used in the remainder of the experiments. Although there are a few settings of the sliding window implementation (i.e., *T*_ud_ = 60 s and *K* = 40) that give a slightly better final accuracy for similar settling times, they require more memory storage (42× more elements) and are also computationally heavier (due to the repredictions of the labels). Lastly, plugging in the hyperparameter values *α* = *β* = 0.9 in (10), which allows converting the forgetting factors *α* and *β* of the recursive implementation to the equivalent pool size parameter *K* of the sliding window implementation, results in *K* = 19 min. This is indeed consistent with the results of the sliding window implementation with *K* = 19, which has a very similar performance (73.6% (st. dev. 13.2%) in 19.4 min (st. dev. 11.5 min)) while requiring much more memory. There is no noticeable benefit from the exponential weighting over the uniform weighting, given that the performance is tested on an asynchronous, independent test set. In Section V-B, we will concurrently test and update on the same data, potentially revealing the benefit of exponential weighting.

### D. Effect of repredictions

The results in Section IV-C show that the single-shot recursive implementation without repredictions of the labels performs on par with the sliding window implementation with repredictions of the labels of previous segments. This suggests that, in the considered time-adaptive context, the iterative repredictions of labels on the current pool of *K* segments have no additional benefit and that the labels hardly change between before and after the relabeling procedure. This is confirmed by computing the total number of labels that changed between before and after the iterative relabeling procedure in all updates before the settling time, i.e., before reaching steady-state performance.

For *T*_ud_ = 60 s, this total number of labels that changed is (on average across subjects and random permutations in Section IV-C) only 0.16 for *K* = 10, 0.70 for *K* = 20, and 1.13 for *K* = 40. This shows that the number of self-corrected labels in the iterative relabeling procedure is minimal, even more so if the pool size *K* is small.

To more closely investigate this dependence of the relabeling on the pool size *K*, we compute the training accuracy (i.e., the percentage of correct labels in the updating set) in the different iterations as a function of the size of the updating set for the batch-mode unsupervised algorithm in [8] using *T*_ud_ = 60 s updating segments and *T*_test_ = 30 s decision segments. Per subject and size of the updating set, ten runs with different randomly selected updating sets of size *K* from the total dataset are performed. Figure 6 then shows the average training accuracy across subjects and runs on Dataset I. From Figure 6, it is clear that the self-correcting behavior on the predicted labels of the first iteration only starts to occur when the updating set contains more than 14–20 min of data. This can be explained from a mathematical point of view as an overfitting effect: when *K* is small, the decoder has enough degrees of freedom to span all initial predictions in the subsequent iterations, leading to an overfitted decoder.

When *K* ≥ 14, there is a clear effect of the second and subsequent iterations (until convergence to the fixed point [8]). This effect, however, seems not to be present in the time-adaptive context. This is explained by considering the initial decoder for each new decoder update when a new segment becomes available. In the batch-mode design, this initial decoder is always a random decoder, whereas in the time-adaptive context, this will only be the case for the first received data segment. In the later updates, the initial decoder is already improved based on past data. Consider the case of *T*_ud_ = 60 s*, K* = 20 in Figure 6. In the time-adaptive case, already 19 updates will have been completed before a full pool of *K* = 20 segments becomes available. After 19 updates, however, the decoder has already substantially improved (see also Figure 3). Therefore, as suggested in Figure 6, the decoder will not exhibit random performance but performance close to the one of the converged decoder in Figure 6. Consequently, the effect of a reprediction on the labels on previous segments in the pool will be similar to the effect of the last iterations (3, 4, 5) in Figure 6, that is, very small. In other words, the initial decoder is then already close to the fixed point of the updating procedure [8].

Given the important trade-off in memory usage and computational complexity, these insignificant improvements do not outweigh the additional required resources.

### E. Validation on an independent dataset

To confirm that the recursive implementation with *T*_ud_ = 60 s and *α* = *β* = 0.9 is a robust choice across subjects and datasets, and that no overfitting of the hyperparameters has occurred on Dataset I, we apply the recursive algorithm on the completely independent Dataset II, again starting from a fully random decoder. Given that Dataset II only contains 50 s trials, we choose *T*_ud_ = 50 s. Furthermore, given that *α* = *β* = 0.9 is equivalent with *K* = 19 according to (10), resulting in a 19 min history when using 60 s segments, this becomes *K* = 22.8 for 50 s segments. Using (10), the equivalent choice for Dataset II becomes *α* = *β* = 0.916.

We test this recursive implementation with *T*_ud_ = 50 s and *α* = *β* = 0.916 on each of the 18 subjects of Dataset II by ten times randomly selecting 40 min as the updating set and the remaining 10 min as test set. The average final accuracy and settling time (on 30 s decision segments) are 76.2% (st. dev. 12.4%) and 18.8 min (st. dev. 10.3 min) (see Figure 5 for the per-subject performances). As this is very similar to the performance obtained in Section IV-C (and even slightly better), it confirms that the chosen specific recursive implementation is a robust choice.

## V. EVALUATION IN TIME-ADAPTIVE CONTEXT

In Section IV, we have tested the proposed time-adaptive unsupervised stimulus reconstruction algorithm asynchronously, i.e., the test set is time-independent from the updating set. While these experiments allowed to investigate the behavior of the proposed method, they do not necessarily reflect a practical use case of the algorithm as in a neuro-steered hearing device application, i.e., while the time-adaptive unsupervised decoder needs to simultaneously update/adapt and provide AAD decisions. In Section V-A, we simulate on Dataset I a situation where electrodes are disconnected, for example, due to movements. In Section V-B, we then evaluate the time-adaptive unsupervised decoder on a third dataset, i.e., while needing to adapt across multiple recording days.

### A. Suddenly disconnecting EEG electrodes

#### 1) Experiment

We compare the fixed supervised decoder with the proposed time-adaptive unsupervised decoder using the selected recursive implementation with *α* = *β* = 0.9 and *T*_ud_ = 60 s from Section IV, when simulating a situation where electrodes are disconnected. Per subject of Dataset I, we randomly permute the 6 min-trials of the first 48 min, to which we add the last 24 min of repetition data, resulting in 72 min of data. The fixed supervised decoder is trained on the first 30 min (i.e., using the available attention labels). Furthermore, also during these first 30 min, the time-adaptive unsupervised decoder has time to update itself starting from a fully random initial decoder, with the autocorrelation matrix and crosscorrelation vector initialized with all zeros.

After these first 30 min of data, we simulate the case where a number of electrodes are disconnected, as could occur in practice, by setting some EEG channels to zero. On these last 42 min of data with disconnected electrodes, the original fixed supervised decoder (trained with all electrodes) is then applied on each *T*_test_ = 30 s decision segment, while the time-adaptive unsupervised decoder keeps on continuously updating per *T*_ud_ = 60 s, and decoding the auditory attention per *T*_test_ = 30 s decision segment. We then compare both decoders after the electrodes are disconnected, i.e., by computing the accuracy across all binary decisions on the last 42 min (thus also taking the settling period of the adaptive decoder after the change into account).

This experiment is performed in two scenarios: when starting from the full high-density 64-channel EEG setup and from a reduced 22-channel subset, where the electrodes are selected corresponding to the mobile 24-channel SMARTING EEG system from mBrainTrain. The latter is added to compare the results with those in Section V-B, where a third dataset is introduced that is recorded using this 24-channel EEG system. The number of disconnected electrodes is varied from 0 to 32 (for the 64-channel case) and from 0 to 11 (for the 22-channel case). Per number of disconnected electrodes, ten random permutations (i.e., of randomly permuting the first eight 6 min-trials and set of disconnected electrodes) are performed.

#### 2) Results

Figure 7a shows the average accuracy across all 16 subjects of Dataset I and the ten random permutations (per subject and number of disconnected electrodes) as a function of the number of disconnected electrodes, starting from the full high-density 64-channel setup and reduced 22-channel setup. When no electrodes are disconnected, the fixed supervised decoder outperforms the time-adaptive unsupervised one with around 4.4% in accuracy in both cases. This difference in accuracy is expected and in line with the batch results obtained in [8]. However, in the 64-channel setup, already when disconnecting three electrodes, the adaptive unsupervised decoder performs better than the fixed supervised one. This effect is already present after two electrodes for the more mobile setup of 22 channels. The difference between both decoders then increases up to 13.9% for the 64-channel setup and 8.3% for the 22-channel setup.

**Figure 7:**
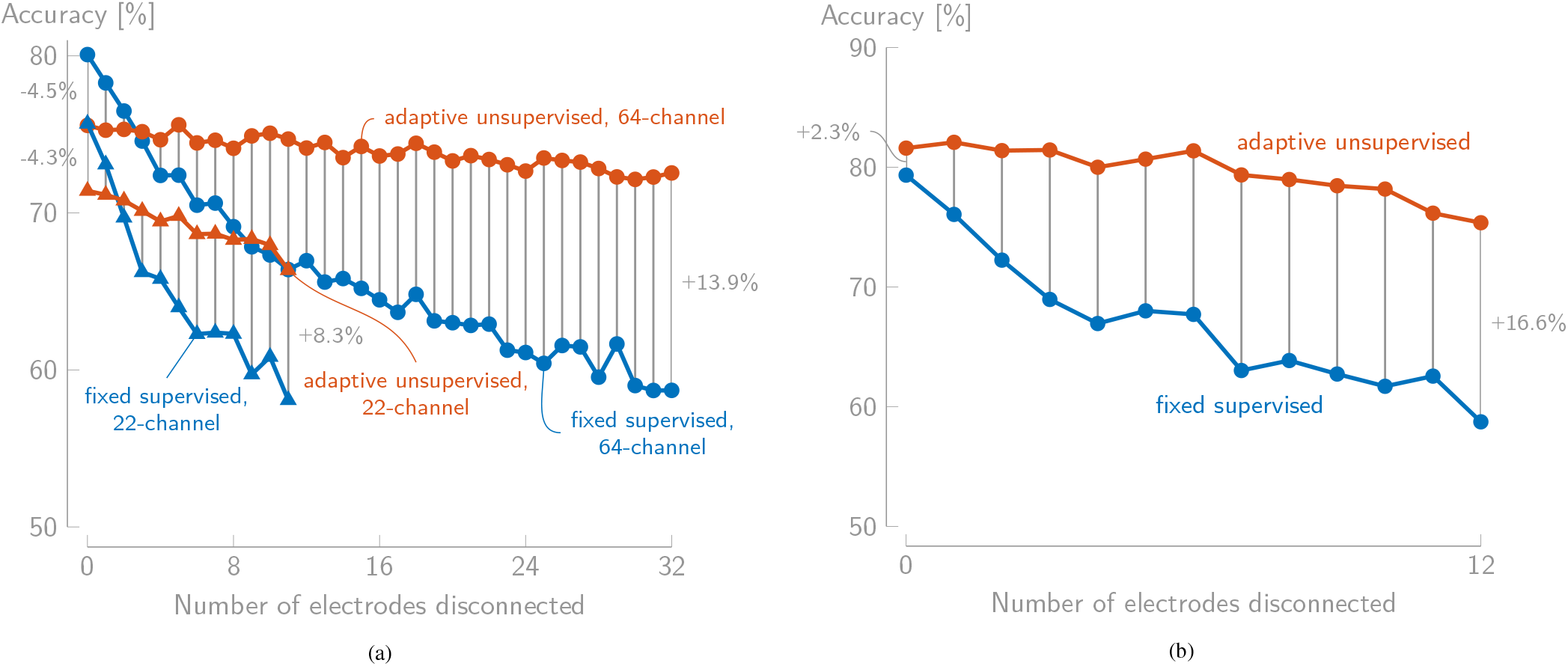
The accuracy on 30 s decision windows of the fixed supervised and time-adaptive unsupervised decoder as a function of the number of disconnected electrodes for (a) Dataset I (electrodes are disconnected after the first 30 min of (training) data) and (b) Dataset III (electrodes are disconnected after the first two (training) sessions). The accuracies are averaged over all 16/2 (Dataset I/Dataset III) subjects and ten runs (per subject and number of disconnected electrodes).

These performance differences seem to be mainly due to the decrease in accuracy of the fixed supervised decoder, which has been trained without taking the disconnected electrodes into account, while the time-adaptive unsupervised decoder remains relatively stable (especially in the 64-channel case). This shows that the latter decoder can effectively adapt to disconnected electrodes, quickly finding an almost equivalent alternative way to decode the attended speech envelope from the reduced set of electrodes. The fact that the adaptive decoder obtains similar performances with only 32 channels compared to 64 channels comes not as a surprise, given that in [18], it was shown that the number of EEG channels could be reduced to around ten without a loss in performance, however, given an optimal channel selection procedure (while here we simulate random electrode disconnections, as would occur in practice).

This experiment clearly shows the added value of the time-adaptive unsupervised approach, effectively and *automatically* adapting to changes in the EEG setup, here simulated by electrodes that are disconnected. Furthermore, we have only simulated one change in the (EEG) setup in an otherwise very controlled experiment, and already obtained a better performance with the time-adaptive approach when two or three electrodes are disconnected. In practice, such changes would occur in combination with other non-stationarities in the data, which is investigated in the next section.

### B. Adaptation across multiple recording days

While in Section V-A, the proposed time-adaptive unsupervised decoder is tested in a more time-adaptive context where electrodes are disconnected, the non-stationarities in the data are still limited to this single change. Furthermore, the EEG data per subject are recorded in one session, with only small breaks in between, and in a very controlled setup. Therefore, in this section, we evaluate the proposed method on a third dataset where the decoder needs to adapt across multiple days of recordings, potentially combined with electrodes that are disconnected.

#### 1) Data and preprocessing

##### a) AAD dataset

We use a third dataset (Dataset III) containing EEG and audio data of two subjects from a longitudinal AAD experiment across multiple days, carried out at the participants’ homes. A two-talker AAD experiment was conducted in eight different sessions that took place on seven different days. In each session, four blocks of 6 min stories are presented to each subject, resulting in a total of 192 min of AAD data per subject. The audio stimuli differed across all sessions. The first two sessions took place on the same day, while the other six sessions took place on different days (see also Figure 8a). The EEG was measured using a 24-channel SMARTING mobile EEG system from mBrainTrain at a 500 Hz sample rate. More detailed information about this dataset can be found in [19].

**Figure 8:**
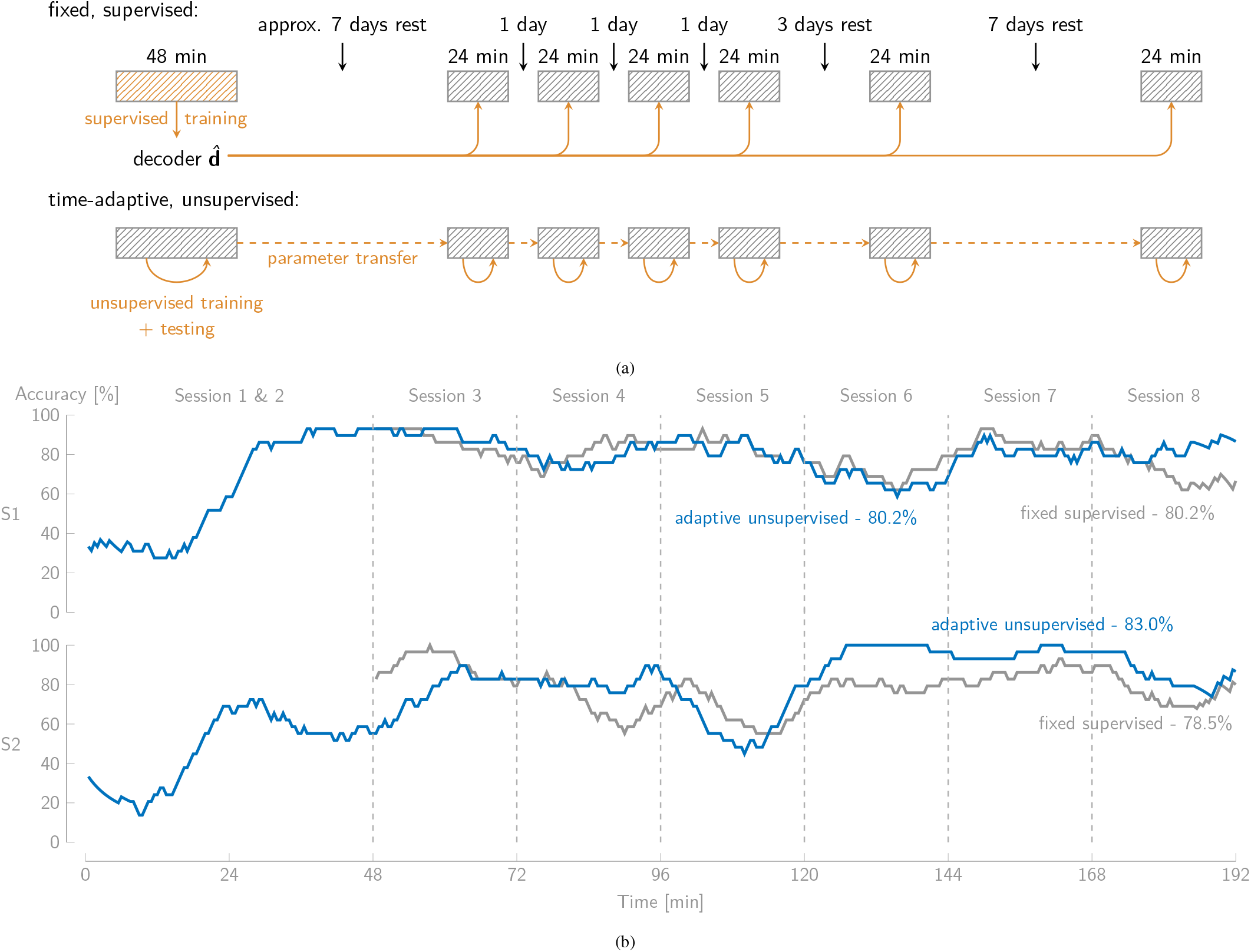
(a) The setup of Dataset III. The fixed decoder is trained in a supervised manner on the first two sessions on the first day and tested on all other sessions on the other days. The proposed time-adaptive unsupervised decoder is initialized with a random decoder and is continuously updated over time and tested/applied on/to each next two decision segments. The autocorrelation matrix and crosscorrelation vector are stored in between sessions (‘parameter transfer’). (b) The smoothed accuracy as a function of time of the fixed supervised and adaptive unsupervised decoder for both subjects of Dataset III. The first 48 min are used as the training set for the fixed supervised decoder, explaining why there is no accuracy there. The final accuracies are computed from Session 3 onward on 30 s decision windows.

While this dataset was initially recorded for the purpose of neurofeedback experiments, it can be used to test the proposed time-adaptive unsupervised decoder as it reflects the practical use case of a neuro-steered hearing device. The algorithm will need to adapt over various days, meaning that there will be changes in, for example, EEG setup, electrode impedances, conditions, speaker and story characteristics, and state of mind of the user, as would all occur in practice. While two subjects are not enough to draw firm (statistical) conclusions, it allows showcasing how the proposed algorithm can be used in a practical, online context.

##### b) Preprocessing

The EEG data and audio envelopes are preprocessed in the same way as in Section IV-A. The only difference is that *L* = 400 ms is chosen as the post-stimulus range of time lags for the decoder, reflecting the choice in [19].

#### 2) Experiment

We compare the fixed supervised decoder with the proposed time-adaptive unsupervised decoder. Again, we only consider the recursive version of the time-adaptive decoder, as it performs similarly to the sliding window version while requiring much less resources. As shown in Figure 8a, the supervised, fixed decoder is trained on the 48 min of data from the first two sessions on the first day, using the information about which speaker is the attended one. This fixed decoder is then applied per *T*_test_ = 30 s decision segment on all other sessions on the other days. As such, this decoder reflects the practical use case where first data of a new neurosteered hearing device user need to be recorded in an a priori calibration session, whereafter the trained decoder is loaded onto the device.

The time-adaptive unsupervised decoder is implemented using the settings as determined in Section IV and is initialized with a fully random decoder, while the autocorrelation matrix and crosscorrelation vector are initialized with all zeros. Per *T*_ud_ = 60 s, the decoder is continuously updated using the recursive implementation with *α* = *β* = 0.9. To fully leverage the time-adaptivity of this decoder, after each update, it is, similarly to Section V-A, applied to the next two *T*_test_ = 30 s decision^2^ segments to make AAD decisions (Figure 8a). The first 48 min of the first two sessions on the first day are used to let the decoder initialize and converge as in Figure 3, starting from a random initial decoder. In between sessions, as one would do in a practical scenario as well, the current autocorrelation matrix **R**_*xx*_ and crosscorrelation vector **r**_*xs*_ are re-used (hence the ‘parameter transfer’ in Figure 8a) and not re-initialized each time from scratch.

Per 30 s decision segment, a decision about the attended speaker is made, resulting in a binary correct/incorrect decision. To provide a comprehensible plot when plotting the accuracy over time, these binary decisions are smoothed using a 29-point moving average (i.e., per segment taking the past and following 7 min into account). To assure a fair comparison between the fixed supervised and time-adaptive unsupervised decoder, the total accuracy is computed as the average over all binary decisions across all but the first 48 min of the first two sessions on the first day.

Lastly, similarly to Section V-A, we evaluate the performance in case one or more electrodes are disconnected by simulating 0 to 12 disconnected electrodes after the first two sessions (see also Section V-A1). Per number of disconnected electrodes, ten random selections of electrodes are performed.

#### 3) Results

Figure 8b shows the smoothed accuracy as a function of time of both the fixed supervised and time-adaptive unsupervised algorithm for both test subjects. As explained before, there is no test accuracy present in the first 48 min for the fixed supervised decoder, as it is trained on those sessions. During those first two sessions, the time-adaptive unsupervised decoder converges after ±25 min, starting from a random initial decoder. This is more or less in line with the results of Section IV-C and IV-E.

The fixed supervised decoder reaches a total accuracy of 80.2% (subject 1) and 78.5% (subject 2), while the proposed time-adaptive unsupervised decoder reaches a total accuracy of 80.2% and 83.0%. The latter thus performs on par with the former for the first subject while outperforming the former with 4.5% for the second subject. Furthermore, our approach does not require an a priori calibration session with the end-user but can be implemented in a plug-and-play fashion on a device. Lastly, more severe changes in the setup and conditions can occur. For example, Figure 7b shows the results of simulating disconnected electrodes. When no electrodes are disconnected, the accuracies are the same as in Figure 8b. However, when one electrode is disconnected, the time-adaptive unsupervised decoder already outperforms the fixed supervised decoder with 6.0%, increasing to 16.6% when 12 electrodes are disconnected. This shows that the proposed method would be able to adapt to such changes, while the fixed supervised decoder only performs worse.

To evaluate whether, besides the favorable memory usage, the recursive implementation also benefits from the exponential weighting compared to the uniform weighting in a sliding window implementation in a time-adaptive context, we test the sliding window implementation with *K* = 19 (i.e., equivalent to *α* = *β* = 0.9) but without repredictions of the labels (to be in line with the recursive implementation which also does not repredict previous labels). The resulting accuracy is 79.9% (subject 1) and 81.6% (subject 2). While this is for both subjects worse than the recursive implementation, we cannot draw firm conclusions about this based on two subjects alone. However, as expected, these results at least suggest that an exponential weighting is favorable compared to a uniform weighting in a time-adaptive context.

Although data from only two subjects are available in this experiment, hampering clear statistical conclusions, the results clearly show the potential of the proposed time-adaptive unsupervised decoder in a practical AAD use case.

## VI. DISCUSSIONS AND CONCLUSIONS

We adapted the offline batch version of the unsupervised AAD stimulus reconstruction algorithm as proposed in [8] to a time-adaptive online version. This allows the decoder to automatically adapt to non-stationarities in the EEG and audio. We have developed both a sliding window implementation with repredictions of previous labels in a finite pool, and a single-shot predict-and-update recursive implementation without repredictions. The latter has the advantage, as it results in similar performances for much less memory usage and computational requirements (Section IV). We have selected the algorithm’s hyperparameters via extensive experiments and validated these on an independent dataset. Furthermore, we explained why there are hardly any changes in labels when using iterative repredictions in this time-adaptive context while this is the case in the batch-mode algorithm presented in [8].

We have also shown the additional benefit of the time-adaptive unsupervised decoder compared to the fixed supervised decoder in a time-adaptive scenario, for example, when simulating electrode disconnections (Section V-A). When electrodes are disconnected, the former starts to clearly outperform the latter. Lastly, the proposed time-adaptive unsupervised decoder outperformed the fixed supervised decoder on a dataset that reflects a practical AAD use case (with testing across multiple sessions on different days; Section V-B). Given that this dataset only contains two subjects, we are careful in drawing firm conclusions. The results, however, clearly show the potential of the proposed method.

As explained in [5], [8], the stimulus reconstruction method does not perform well enough on short decision segment lengths for the online AAD application. In Section V-B, there were no switches in auditory attention present, such that a high accuracy could be obtained with 30 s decision segments. While this reduces the relevance of the proposed algorithm as the ‘decision-maker’ in AAD, it is an excellent candidate to provide reliable labels of the auditory attention to update a faster algorithm (such as the common spatial pattern filtering algorithm [20]) in an unsupervised manner but only at low speeds. As an automatic ‘labeler’ to inform another (supervised) AAD algorithm, the speed at which labels are produced is less restricted.

To conclude, the proposed time-adaptive unsupervised stimulus reconstruction method is an important step forward to the online application of AAD in neuro-steered hearing devices.

## Footnotes

1 The speech envelope can be defined and extracted in different ways [9]. In this paper, we use the speech envelope extract procedure based on a gammatone filterbank and power-law compression as proposed in [9].

2 Two 30 s decision segments as the decoder is only being updated every 60 s.

## Notes

This work is supported by an Aspirant Grant from the Research Foundation Flanders (FWO) (for S. Geirnaert - 1136219N), FWO project nr. G0A4918N, the European Research Council (ERC) under the European Union’s Horizon 2020 Research and Innovation Programme (grant agreement No 802895), and the Flemish Government (AI Research Program). The scientific responsibility is assumed by its authors.

### Competing Interest Statement

The authors have declared no competing interest.

